# Phenotypic variation in divergent mitochondrial haplotypes in migratory three-spined stickleback

**DOI:** 10.64898/2026.11.04.764197

**Authors:** Emma Shepherd, Megan Barnes, Henry Lewis, Ben Fisher, Andrew MacColl

## Abstract

Mitochondria are essential components in life-sustaining organismal processes. The mitochondrial genome can be influenced by selection, and divergent mitochondrial haplotypes have substantial potential to affect interspecies variation in life-history and physiology, yet little is known about whole organism phenotypic changes that might accompany mitochondrial genomic divergence. Three-spined stickleback on the Scottish island of North Uist provide a model to study this as their populations exhibit a well-mixed nuclear genome, yet two highly diverged mitochondrial haplotypes (‘trans-Atlantic’ and ‘European’). Three-spined stickleback were sampled from six lochs to investigate associations between mitochondrial haplotype and phenotypic traits including organ size, body size, and ovule volume/clutch size, as well as citrate synthase enzyme activity. Results showed that mitochondrial haplotype has an influence on male gonadosomatic index (GSI) and potentially citrate synthase (CS) enzyme activity. Differences in male GSI may be explained by the timing of male spermatogenesis or potentially a need for larger testes in European haplotype fish. While no other phenotypic trait showed differences between mitochondrial haplotypes, there were many differences between sexes and populations e.g. in body size and organ/muscle size.

## Introduction

Mitochondria are the main locations of energy production within an organism and contribute to controlling the ways in which that energy is used. ATP generated within the mitochondria is used for life sustaining organismal processes like macromolecule synthesis, muscle contraction, and thermogenesis (Bonora et al. 2012). The mitochondrion also has its own genome that can accumulate mutations that may influence mitochondrial function (Wolff et al. 2016) and potentially be under selection. Divergent mitochondrial DNA haplotypes have substantial potential to contribute to the life-history and physiological variation seen between species (Scott & Milsom, 2011, Kurbalija Novicic et al. 2015, Dowling & Wolff 2023), but surprisingly little is known about their contribution to phenotypic variation within natural populations. This is probably because strongly divergent mitochondrial haplotypes seldom segregate within populations or against a constant nuclear genetic background. This makes it difficult to separate the main effects of mitochondrial versus nuclear genomic variation on phenotype. This is further complicated by the possibility of mito-nuclear interactions (incompatibilities) (Healy & Burton 2020, Torres-Gonzalez & Makova 2022).

Some studies have studied the effects of mitochondrial variation on life-history and physiology in laboratory settings while controlling for the nuclear genome (Latorre-Pellicer et al. 2016, Dowling & Wolff 2023, Rand et al. 2006). These studies usually use experimental ‘conplastic’ strains in which organisms with different mitochondrial haplotypes are compared against the same nuclear background. This is achieved by experimental manipulations involving repeated backcrossing or direct transfer of genetic material between ovules. Conplastic studies are an effective way to look at traits that are directly affected by the mitochondrial genome and have often shown large mitochondrial effects on physiology and life-history (Latorre-Pellicer et al. 2016, Camus et al. 2017) but are rare in non-model organisms. They also do not always completely escape the possibility of mito-nuclear incompatibility, since sometimes only a single nuclear background is used.

A small number of studies have demonstrated substantial effects of mitochondrial variation on phenotypes in natural populations, but these are largely confined to *Drosophila* (Dowling & Wolff 2023, Ballard et al. 2007) and vertebrate examples are lacking. Here we focus on the three-spined stickleback (*Gasterosteus aculeatus*) (“stickleback”) as an ideal model for studying the effects of mitochondrial variation on life-history and physiology of vertebrates in the wild (Beck et al. 2022). We take advantage of an unusual circumstance on the island of North Uist, in the Scottish Western Isles, where deeply divergent “European” and “trans-Atlantic” mitochondrial lineages (Mäkinen & Merilä 2007) segregate at roughly equal frequencies in migratory stickleback populations (Dean et al. 2019). It is likely that these lineages diverged in allopatric glacial refugia on opposite sides of the Atlantic and have come into secondary contact on the eastern margins of the Atlantic. Whole genome sequencing reveals no nuclear differences between the lineages within migratory populations (Barnes et al. *in review, PNAS*). Here we compare various phenotypic traits of wild-caught migratory stickleback from multiple Uist populations to ascertain whether mitochondrial genotype might be contributing to physiology and life-history variation.

## Methods

All three-spined stickleback sampled in this study were collected on the island of North Uist in the Scottish Western Isles (57° 34’ 13.4” N; 7° 16’ 52.3” W). Adult migratory stickleback were collected from six different brackish or freshwater lochs (Table 1, Figure 1). These lochs were chosen for sampling due to the prevalence of the migratory stickleback ecotype. All sampling was completed between the 21^st^ of April and the 22^nd^ of May 2025. Migratory stickleback on North Uist, in contrast to resident fresh- and saltwater populations are ubiquitously relatively large, with complete sets of bony lateral plates, large spines and are generally of a silvery colouration (MacColl et al. 2013, Dean et al. 2019, Dean et al. 2024)

**Figure 1.**
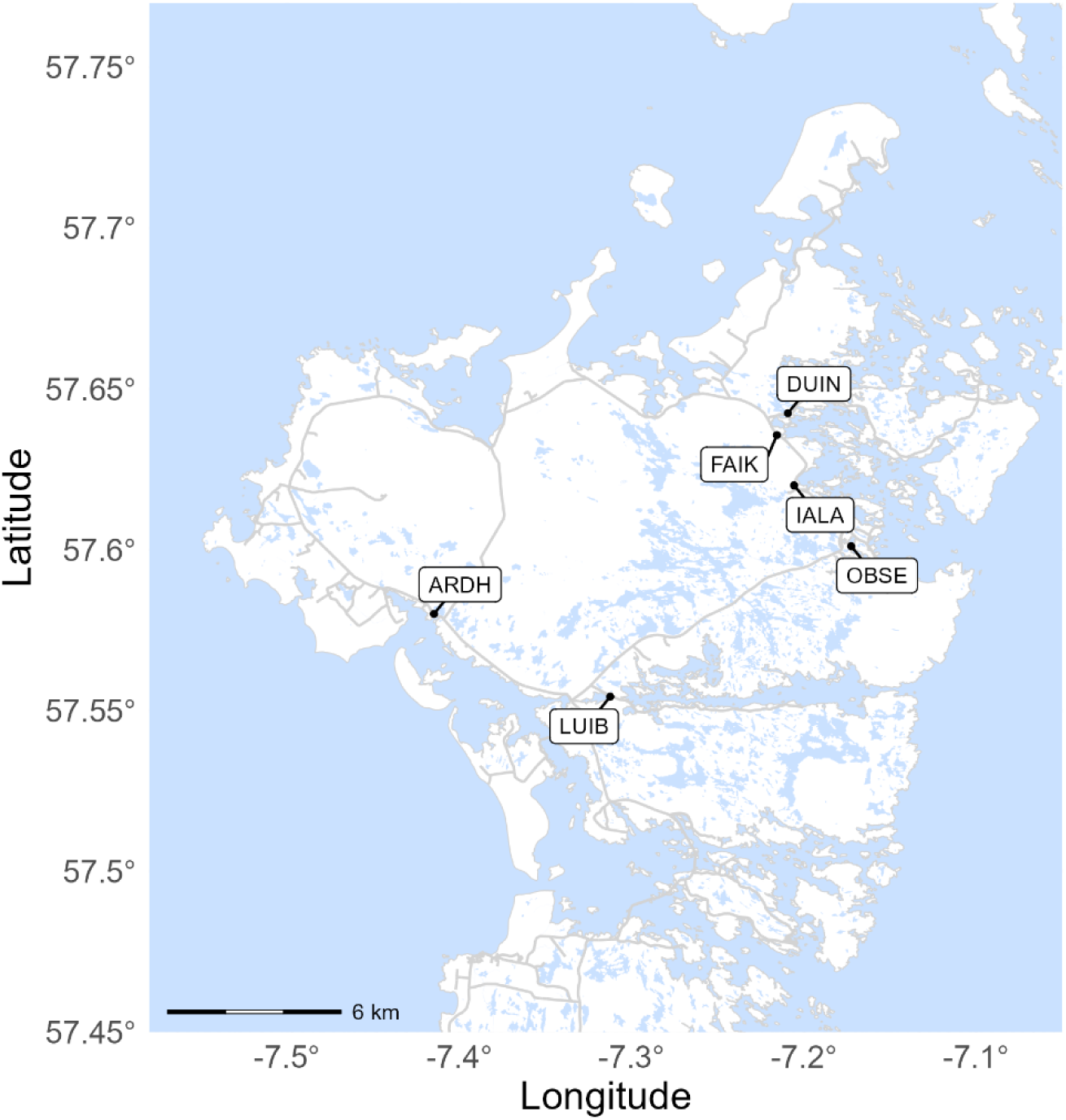
Map of North Uist showing the precise locations of the six sampling sites ARDH, LUIB, DUIN, OBSE, FAIK, and IALA in longitude and latitude.

**Table 1.** Locations of sampled lochs on the Scottish island of North Uist, with the associated coordinates and codes used. Sample size is split by males and females.

| Loch | Code | Coordinates | Sample size (n) |
|---|---|---|---|
| Ardheisker | ARDH | 57° 34' 47" N, 7° 24' 51" W | 48 females<br>21 males |
| Clachan an Luib | LUIB | 57° 33' 14" N, 7° 18' 44" W | 43 females<br>20 males |
| Loch an Duin | DUIN | 57° 38' 32" N, 7° 12' 30" W | 9 females<br>6 males |
| Ob nan Stearnain | OBSE | 57° 36' 4" N, 7° 10' 21" W | 9 females<br>4 males |
| Fairy Knoll | FAIK | 57° 38' 7" N, 7° 12′ 54" W | 5 females<br>5 males |
| Ialadh | IALA | 57° 37' 12" N, 7° 12′ 20" W | 5 females<br>5 males |

The stickleback used for this study were collected using minnow traps (Gee traps, Dynamic Aqua, Vancouver) placed in various parts of each loch. Traps were placed in suitable locations and collected after 24 to 72 hours. Samples of captured stickleback were taken to a temporary field laboratory for phenotyping. This study was approved by the University of Nottingham Animal Welfare and Ethical Review Body, and no permit is required in Scotland for the collection of stickleback. The Code of Practice for the Housing and Care of Animals Bred, Supplied, or Used for Scientific Purposes was followed. Fish were euthanized with tricaine methane sulphonate (MS-222) followed by destruction of the brain to confirm death, as required by the UK Home Office, under project license PP5421721. Dissections were carried out immediately following euthanasia. Fish were weighed to the nearest 0.0001g and standard-length measurements were taken from the tip of the snout to the end of the caudal peduncle. Both of the pectoral fins and the caudal fin were removed and stored in 80% ethanol for later DNA extraction. The left pectoral muscles and liver were dissected out of each fish and weighed. Livers were frozen at −20° C for the citrate synthase assay. The sex of fish was established from inspection of the gonads. Our sample consisted of migratory individuals that had recently returned from the sea. Fish were assumed to be one year old as length was unimodally distributed and previous otolith analysis of migratory fish from migratory North Uist populations showed that individuals were the same age (Barnes et al. 2025). Ovules were extracted from fully gravid fish as a whole clutch and were counted, before 10 ovules from each clutch were measured using a microscope and graticule. Ovule length was recorded as the longest axis through the ovule and ovule width as the greatest dimension perpendicular to length. To determine ovule volume, the equation (4/3)(*πAB*^2^) was used where A and B are the half-axis values for length and width, respectively.

Fin clips were prepared for DNA extraction by taking two fins from each sample and cutting them up into smaller pieces. DNA extractions were carried out using the DNeasy 96 Blood and Tissue Kit from QIAGEN. The protocol provided with this kit was followed, with one adjustment made by using only 100µL of buffer AE in the final step instead of 200µL. This was done to create more concentrated DNA samples. Mitochondrial haplotypes were determined using a restriction assay designed by Barnes et al. (2026). This assay used primers to amplify ND4 and ND5 regions of the mitochondrial genome and used HindIII or PstI restriction enzymes to cut DNA based on lineage. In the ND4 region, HindIII cuts a segment of DNA pertaining to the Eu lineage. In the ND5 region, PstI cuts a segment of DNA pertaining to the At lineage. These regions each contain a fixed synonymous substitution between Eu and At haplotypes. Both regions should provide the same information for each sample, eliminating user error.

One PCR reaction required 10µl of Accustart, 7µL of nuclease free water, 1µL of forward primer, and 1µL of reverse primer (being careful to keep the ND4 and ND5 primers separate from each other). 1µL of DNA was added to each ND4 PCR tube and respective ND5 PCR tube (using 2µL total of DNA per extracted fin clip sample). All 48 PCR tubes were then loaded into the G-Storm GS1 thermocycler to run for two hours. PCR followed this cycle: initial denaturation at 94° C for three minutes, then denaturation at 94° C for 30 seconds, annealing at 60° C for 30 seconds, extension at 72° C for 1 minute, with this whole process repeating 36 times. A final extension then occurred at 72° C for five minutes. A 1.5% agarose gel was created to ensure successful amplification. 6µL of each PCR sample ran through the gel at 100V for 25 minutes, along with 4µL of a 100bp ladder (NEB). DNA was then visualized on an iBright CL750 Imaging System (ThermoFisher Scientific) to confirm amplification

All successfully amplified samples were then mixed with 0.8µL of restriction enzyme and 1.5µL of its associated 10X restriction enzyme buffer. For all ND4 samples, HindIII restriction enzyme and r2.1 10X restriction enzyme buffer were added. For all ND5 samples, PstI restriction enzyme and r3.1 10X restriction enzyme buffer were added. These PCR samples were then put back into the G-Storm GS1 thermocycler for incubation at 37° C for one hour. After incubation, 6µL of each PCR sample was run on a 2% agarose gel at 100V for 25 minutes along with 4µL of a 100bp ladder (NEB). At this point, mitochondrial lineages could be visualized using the iBright CL750 Imaging System (ThermoFisher Scientific).

Abcam’s Citrate Synthase Assay Kit (ab239712) was used to measure citrate synthase activity in frozen liver samples from ARDH, OBSE and LUIB. All liver samples were cut and weighed to be as close to 10mg as possible. Following the protocol, standard dilutions, positive control, and background reaction mixtures were also created and used. Immediately after adding reaction mixtures, samples were put into the Epoch Microplate Reader (BioTek Instruments, Inc) for 30 minutes at 25° C and 412nm. Results were captured every minute for 30 minutes and were exported as an excel file by Gen5 Microplate Reader and Imager Software (version 3.10, BioTek Instruments, Inc) for data analysis.

Citrate synthase enzyme activity was determined using the equation:

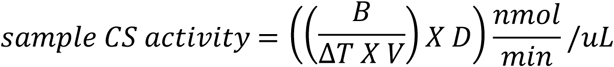

where B is the nanomoles of S-H group from the Standard Curve, △T is the reaction time, V is the sample volume added to each reaction well (10µL), and D is the sample dilution factor (10). B and △T were determined mathematically by plotting time over the optical density of the standards and using the line of best fit (y-0.037B). The value used for △T (10 minutes) was taken from the linear section of the plot. B could then be calculated by subtracting the optical density values at the beginning and end of △T and substituted into the line of best fit equation (Abcam 2023). All citrate synthase values were divided by the weight of the original liver tissue used in grams.

All statistical analyses were conducted using R (version R 4.5.0). Linear models were created using R package “tidyverse” (version 2.0.0, Wickham, 2019). Generalized linear models (GLMs) were created using “stats” (version 4.5.0, R Core Team, 2025). Specific factors used in each model are described in the supplementary material. Explanatory variables included mitochondrial haplotype, population, and sex, as appropriate for each model. Sampling date was initially added as a covariate but was excluded from the final models where this showed no significance (p-value > 0.05). Where relevant (ie. GSI), date of collection was retained in the model. Additionally, body size was accounted for in analyses of ASI, HSI, GSI, liver weight per gram, and pectoral muscle weight per gram as these indices implicitly include body weight as a measure. ANOVA Type 1 sums of squares (GLMs) or ANOVA Type 3 sums of squares (LMMs, using Satterthwaite’s method for degrees of freedom) were run on models to determine significance where appropriate.

Female Gonadosomatic Index (GSI) data were log transformed. One individual in the male GSI dataset was removed as the recorded gonad weight was unclear. For clutch size analysis, the average ovule volume for each clutch was used. Two samples were removed from the citrate synthase analysis, as the kinetics machine did not read properly, producing a negative and zero result.

## Results

We phenotyped and genotyped stickleback from six populations. Sample sizes of some location-sex combinations were small, but the total sample size (n=180) allowed balanced comparisons of mitochondrial genotypes (136 At, 44 Eu) and sexes (119 females, 61 males). The locations and numbers of stickleback sampled in this study can be found in Table 1. The proportion of the At mitochondrial haplotype in each population can be found in Fig. 2 and ranged from 0.20 (IALA) to 0.90 (FAIK).

**Figure 2.**
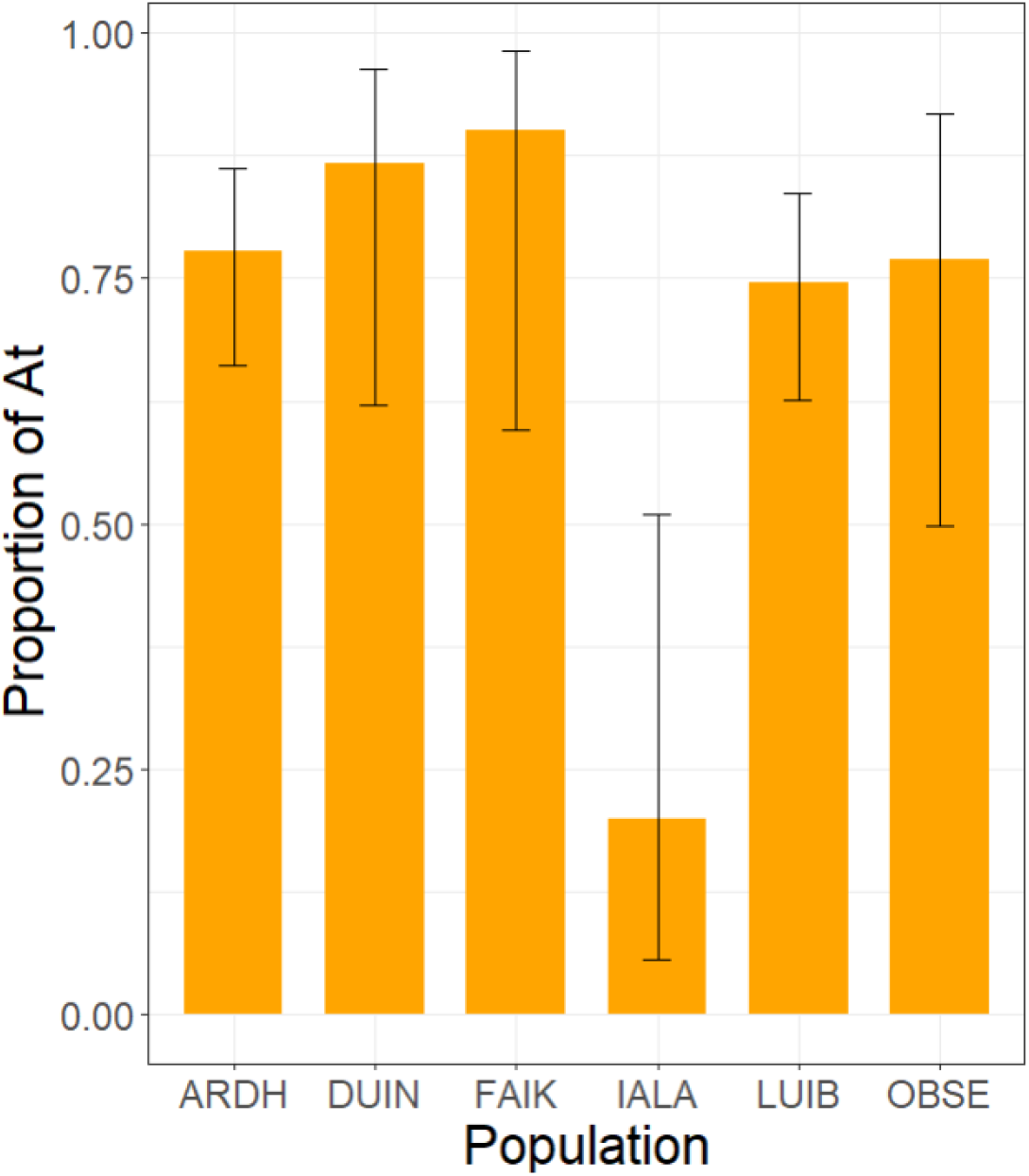
The proportion of the At mitochondrial haplotype present in each population (ARDH, DUIN, FAIK, IALA, LUIB, OBSE). Orange bars highlight the proportion, while black lines represent the 95% binomial confidence intervals.

While there were no significant differences in body size between mitochondrial haplotypes, the stickleback in this study varied in their overall body size between sexes and populations. On average, female stickleback were larger: they were longer and weighed more than their male counterparts. DUIN and FAIK fish were longest while those from ARDH were shortest (Fig. 3, Table 2).

**Figure 3.**
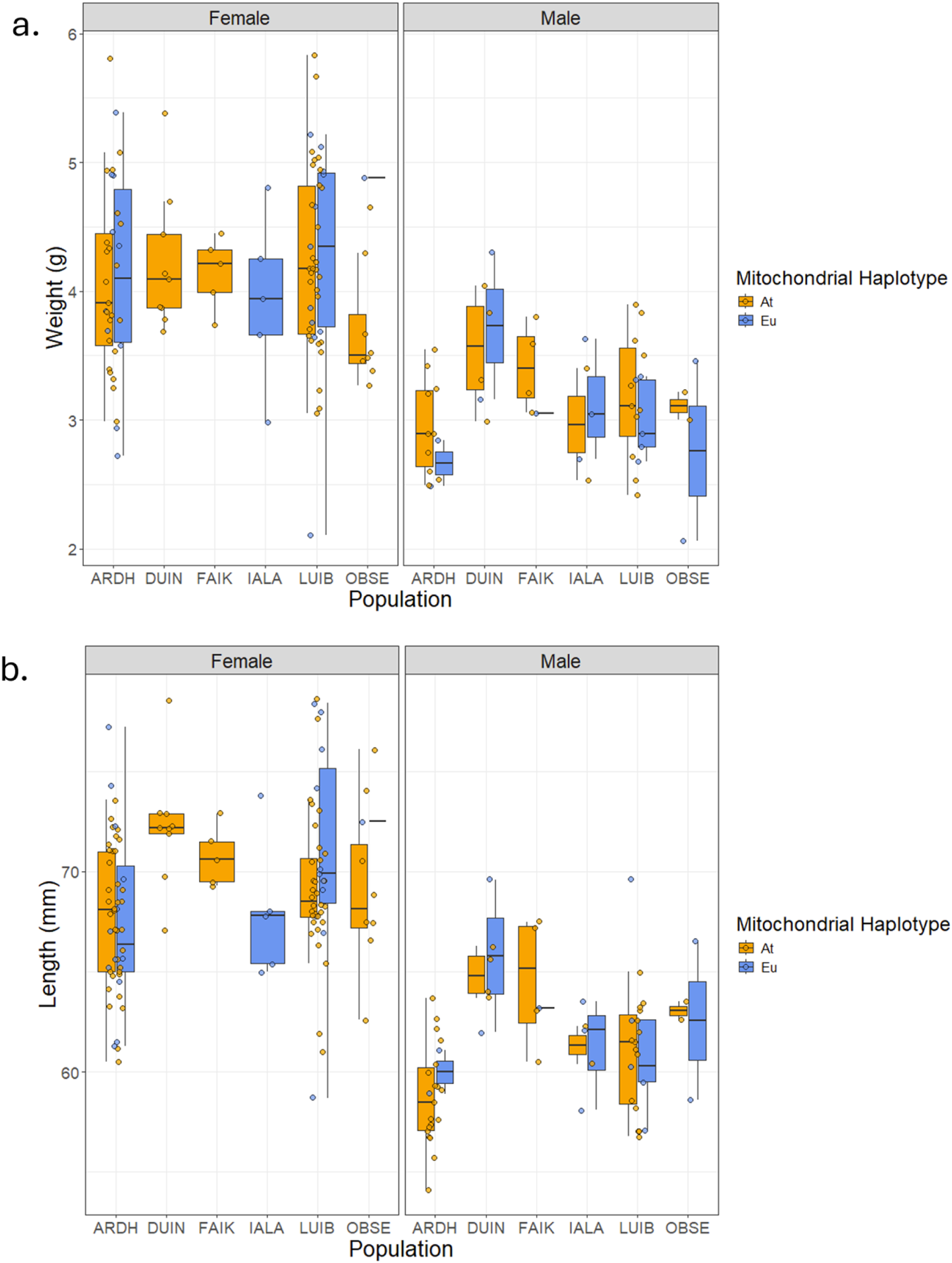
Variation in body size of migratory three-spined stickleback from North Uist, between mitochondrial haplotypes, sexes, and populations (ARDH, DUIN, FAIK, IALA, LUIB, OBSE). a) Weight in grams (female, male). b) Length in millimeters (female, male). Mitochondrial haplotypes are designated for each population as orange for At and blue for Eu. 25 and 75 quartile ranges are shown as vertical black lines. Horizontal black lines on each box plot represent the median. Each individual point represents one adult stickleback sampled.

**Table 2:** Model results^1^ of the variation in phenotypic traits between mitochondrial haplotypes, populations and sexes. Significant p-values are shown in bold.

| **Trait** | **Mitochondrial haplotype** | **Population** | **Sex**<sup>2</sup> | **Model used** |
|---|---|---|---|---|
| **Fish weight (g)** | P-value: 0.9951<br>F-value: 0.0000<br>Mean sq: 0.000<br>Sum sq: 0.000<br>Resid. Df: 142<br>Df: 1 | P-value: 0.2231<br>F-value: 1.4131<br>Mean sq: 0.571<br>Sum sq: 2.857<br>Resid. Df: 142<br>Df: 5 | **P-value: 1.388e-15 (\*\*\*)**<br>F-value: 80.8592<br>Mean sq: 32.695<br>Sum sq: 32.695<br>Resid. Df: 142<br>Df: 1 | Linear<sup>3</sup> |
| **Standard length (mm)** | P-value: 0.2023<br>F-value: 1.6381<br>Mean sq: 20.24<br>Sum sq: 20.24<br>Resid. Df: 172<br>Df: 1 | **P-value: 4.616e-6 (\*\*\*)**<br>F-value: 7.1008<br>Mean sq: 87.73<br>Sum sq: 438.66<br>Resid. Df: 172<br>Df: 5 | **P-value: <2.2e-16 (\*\*\*)**<br>F-Value: 204.6032<br>Mean sq: 2527.89<br>Sum sq: 2527.89<br>Resid. Df: 172<br>Df: 1 | Linear<sup>3</sup> |
| **Gonadosomatic index (GSI), male** | **P-value: 0.001181 (\*\*)**<br>F-value: 12.0724<br>Mean sq: 1.1876e-5<br>Sum sq: 1.1876e-5<br>Resid. Df: 38<br>Df: 1 | P-value: 0. 270948<br>F-value: 1.2724<br>Mean sq: 1.2518e-6<br>Sum sq: 6.2590e-6<br>Resid. Df: 38<br>Df: 5 | **Date of Collection<sup>2</sup>**<br>P-value: 0.245129<br>F-value: 1.3616<br>Mean sq: 1.3395e-6<br>Sum sq: 1.3390e-6<br>Resid. Df: 38<br>Df: 1 | Linear<sup>3</sup> |
| **Gonadosomatic index (GSI), female** | P-value: 0.91185<br>F-value: 0.012<br>Mean sq: 0.00203<br>Sum sq: 0.0020<br>Resid. Df: 89<br>Df: 1 | **P-value: 0.03131(\*)**<br>F-value: 2.5847<br>Mean sq: 0.42508<br>Sum sq: 2.1254<br>Resid. Df: 89<br>Df: 5 | **Date of Collection<sup>2</sup>**<br>P-value: 0.36127<br>F-value: 0.8421<br>Mean sq: 0.13849<br>Sum sq: 0.1385<br>Resid. Df: 89<br>Df: 1 | Linear<sup>3</sup> |
| **Hepatosomatic Index (HSI)** | P-value: 0.464<br>F-value: 0.5392<br>Mean sq: 0.0000598<br>Sum sq: 0.0000598<br>Resid. Df: 140<br>Df: 1 | **P-value: 9.369e-5 (\*\*\*)**<br>F-value: 5.6232<br>Mean sq: 0.0006237<br>Sum sq: 0.0031183<br>Resid. Df: 140<br>Df: 5 | **P-value: <2.2e-16 (\*\*\*)**<br>F-Value: 237.0654<br>Mean sq: 0.0262924<br>Sum sq: 0.0262924<br>Resid. Df: 140<br>Df: 1 | Linear<sup>3</sup> |
| **Liver weight per gram** | P-value: 0.6074442<br>F-value: 0.2651<br>Mean sq: 0.0000245<br>Sum sq: 0.0000245<br>Resid. Df: 140<br>Df: 1 | **P-value: 0.0001115 (\*\*\*)**<br>F-value: 5.5305<br>Mean sq: 0.0005120<br>Sum sq: 0.0025601<br>Resid. Df: 140<br>Df: 5 | **P-value: <2.2e-16 (\*\*\*)**<br>F-value: 174.6949<br>Mean sq: 0.0161733<br>Sum sq: 0.0161733<br>Resid. Df: 140<br>Df: 1 | Linear<sup>3</sup> |
| **Pectoral muscle weight per gram** | P-value: 0.8002<br>F-value: 0.0643<br>Mean sq: 0.00000037<br>Sum sq: 0.00000037<br>Resid. Df: 140<br>Df: 1 | P-value: 0.5833<br>F-value: 0.7556<br>Mean sq: 0.00000438<br>Sum sq: 0.00002190<br>Resid. Df: 140<br>Df: 5 | **P-value: <2.2e-16 (\*\*\*)**<br>F-value: 120.6506<br>Mean sq: 0.00069941<br>Sum sq: 0.00069941<br>Resid. Df: 140<br>Df: 1 | Linear<sup>3</sup> |
| **Fat presence** | P-value: 0.9878192<br>Deviance: 0.000<br>Resid. Dev: 151.58<br>Resid. Df: 140<br>Df: 1 | **P-value: 2.36e-7 (\*\*\*)**<br>Deviance: 39.014<br>Resid. Dev: 151.58<br>Resid. Df: 141<br>Df: 5 | **P-value: 0.0002725 (\*\*\*)**<br>Deviance: 13.251<br>Resid. Dev: 190.59<br>Resid Df: 146<br>Df: 1 | Generalized Linear, family: Binomial<sup>4</sup> |
| **Fat weight (g)** | P-value: 0.7993<br>F-value: 0.0649<br>Resid. Df: 140<br>Df: 1 | **P-value: 2.793e-6 (\*\*\*)**<br>F-value: 7.5196<br>Resid. Df: 140<br>Df: 5 | **P-value: 5.562e-5 (\*\*\*)**<br>F-value: 17.2909<br>Resid. Df: 140<br>Df: 1 | Linear<sup>3</sup> |
| **Adiposomatic index (ASI)** | P-value: 0.76719<br>F-value: 0.0880<br>Mean sq: 0.0275<br>Sum sq: 0.027<br>Resid. Df: 139<br>Df: 1 | P-value: 0.09505<br>F-value: 1.9182<br>Mean sq: 0.5987<br>Sum sq: 2.993<br>Resid. Df: 139<br>Df: 5 | **P-value: 1.671e-7 (\*\*\*)**<br>F-value: 30.3824<br>Mean sq: 9.482<br>Sum sq: 9.483<br>Resid. Df: 139<br>Df: 1 | Linear<sup>3</sup> |

<sup>1</sup> The order of factors used in each model is not reflected by the order in which they appear on the table.
<sup>2</sup> For GSI models, sex was not run as a factor as models were split based on sex. Date of collection was considered when modelling GSI as fish were caught on different days of the breeding season depending on the loch sampled.
<sup>3</sup> R package “tidyverse” version 2.0.0, Wickham, 2019
<sup>4</sup> R package “stats” version 4.5.0, R Core Team, 2025

There were no significant differences between mitochondrial haplotypes in almost any of the organ and tissue traits measured (p > 0.05, Table 2, Figure 4). The exception to this was male gonadosomatic index (GSI), with the Eu haplotype showing higher GSI than the At haplotype (F_1,38_ = 12.07, p = 0.0013, Table 2, Fig. 5). This pattern was consistent across all six populations.

**Figure 4.**
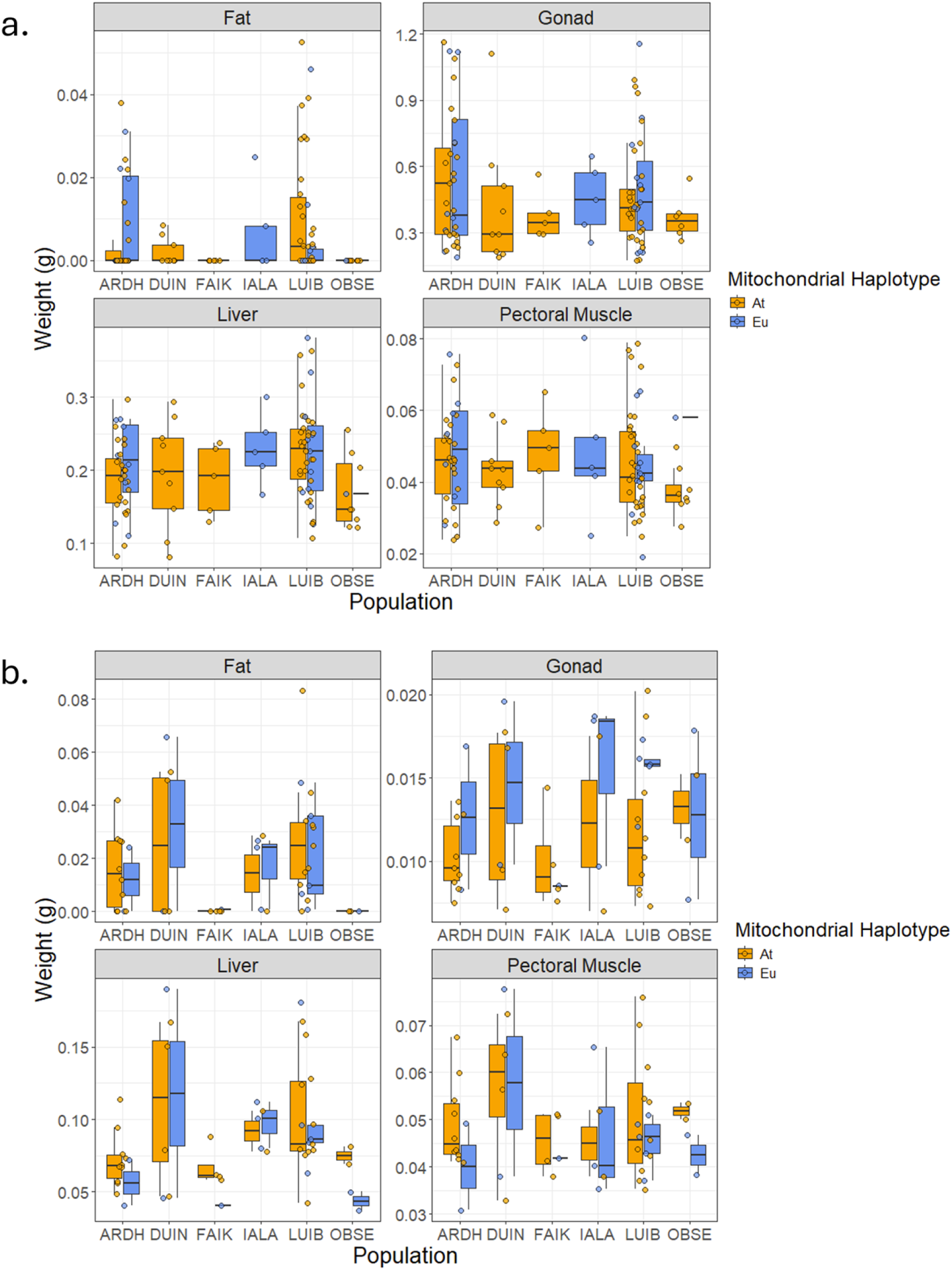
Variation in organ and tissue weights of migratory three-spined stickleback from North Uist, between mitochondrial haplotypes, sexes and populations (ARDH, DUIN, FAIK, IALA, LUIB, OBSE). a) Females (fat, gonad, liver, and pectoral muscle weight in grams). b) Males (fat, gonad, liver, and pectoral muscle weight in grams). Mitochondrial haplotypes are designated for each population as orange for At and blue for Eu. 25 and 75 quartile ranges are shown as vertical black lines with points representing outliers. Horizontal black lines on each box plot represent the median. Each individual point represents one adult stickleback sampled.

**Figure 5.**
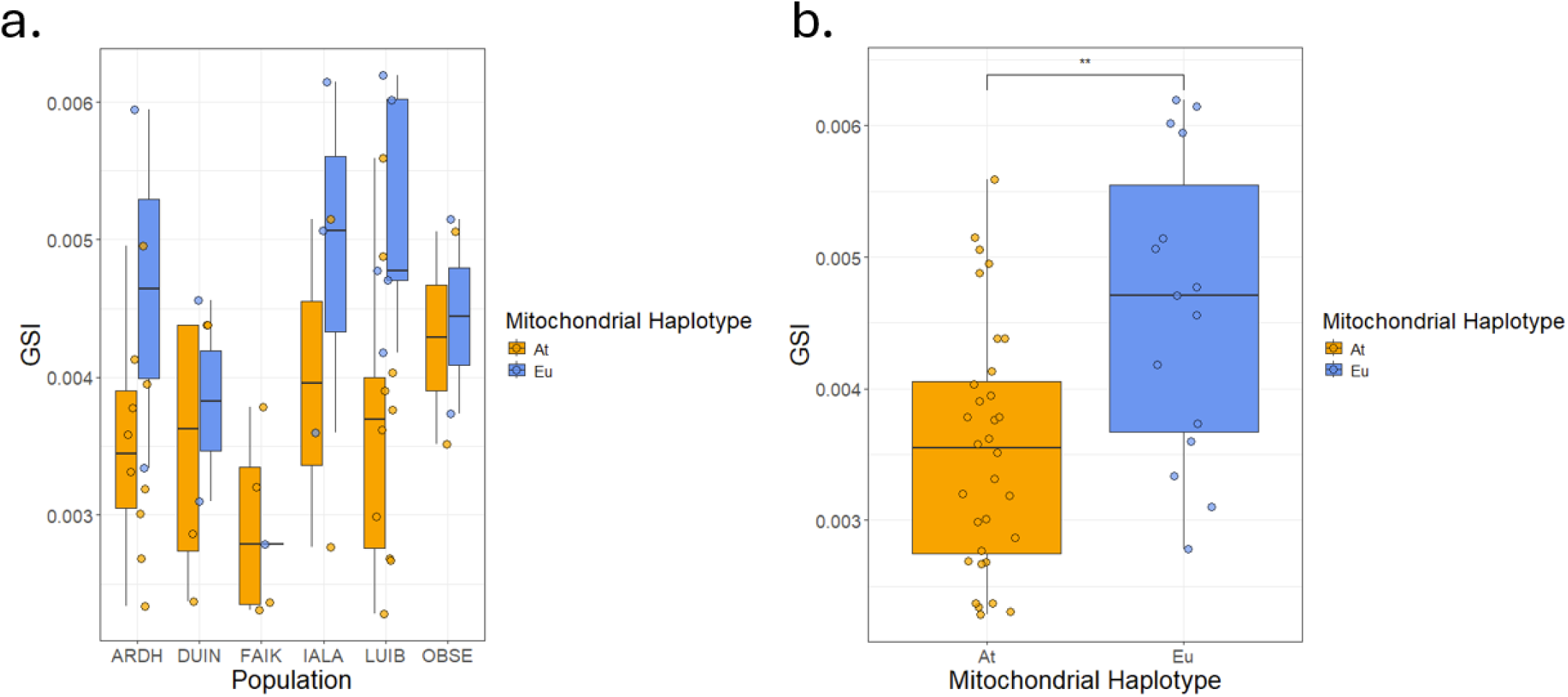
Variation in male GSI measurements. Eu fish had significantly higher GSI compared to At fish. This trend can be visualized across all populations (ARDH, DUIN, FAIK, IALA, LUIB, OBSE). a) GSI differences by population. b) Overall GSI differences between mitochondrial haplotypes. Mitochondrial haplotypes are designated for each population as orange for At and blue for Eu. 25 and 75 quartile ranges are shown as vertical black lines. Horizontal black lines on each box plot represent the median. Each individual point represents one adult stickleback sampled.

There were significant differences in the traits measured between the sexes. Females had heavier livers (relative to overall body size), while males had bigger pectoral muscles and more fat than females (p < 0.05, Table 2, Fig. 4). Differences among populations were also observed. HSI, liver weight per gram, fat presence, fat weight, and female GSI all varied amongst populations (p < 0.05, Table 2, Fig 4, Fig. 6). FAIK and LUIB were the most different from each other across all liver and fat related traits, with LUIB and IALA specifically having the biggest livers. In analyzing these data, we noticed that the longest fish (FAIK) did not have the biggest livers, so we modeled raw liver weight as a function of fish length, population, and their interaction to test for differences in liver allometry. The interaction was not significant, implying that liver size/growth scaled similarly between populations (F_1, 134_=0.44, p = 0.82).

**Figure 6.**
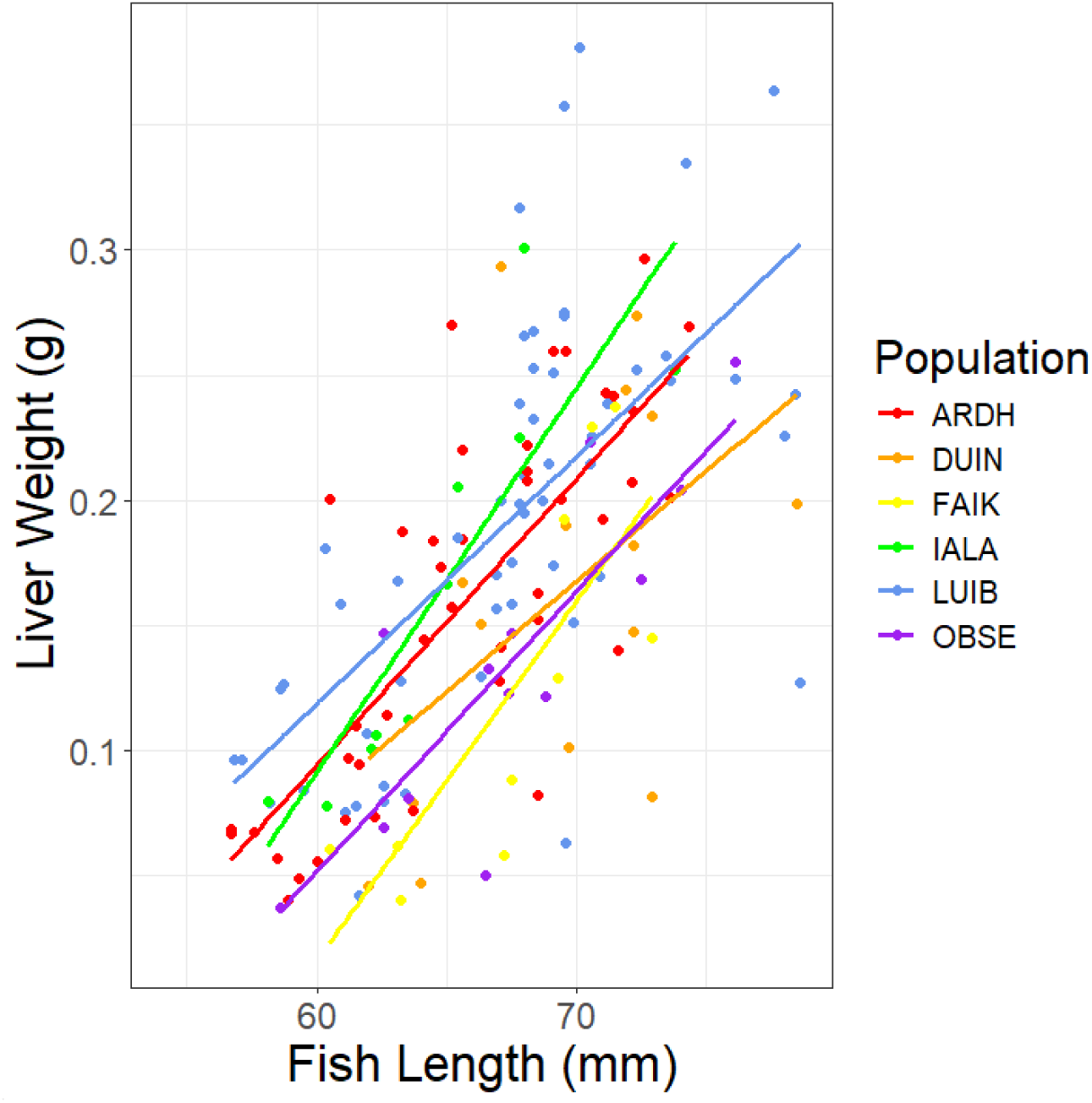
Variation between populations shows the relationship between liver weight and fish length. Populations (ARDH, DUIN, FAIK, IALA, LUIB, OBSE) are designated by the various colors. The lines of best fit indicate the change in liver size with increasing fish length, with each point showing an individual.

Ovules were extracted as a whole clutch from 15 female stickleback (12 from ARDH, one from DUIN, one from IALA, and one from LUIB) and were counted. Clutch size increased with female length but there was no significant difference in clutch size between mitochondrial haplotypes although the sample size for this analysis was small (GLM with Type 1 sums of squares: standard length, F_1,12_ = 6.94, p = 0.022; mitochondrial haplotype, F_1,12_= 0.03, p = 0.86). Ten ovules per stickleback were isolated and used for ovule volume measurements. There was no significant difference in ovule volume between mitochondrial haplotypes or with fish length but, again, sample size was small (generalized linear mixed model with normal errors, stickleback ID as the random effect, Type 3 sums of squares: fish length F_1,11.998_= 2.62, p = 0.13; mitochondrial haplotype, F_1,11.998_ = 1.1520, p = 0.3042).

Livers from 40 wild caught stickleback (21 from ARDH, 12 from OBSE and seven from LUIB) were used to analyze citrate synthase enzyme activity. Citrate synthase activity did not vary with liver weight or mitochondrial haplotypes although it is worth noting that the means for Eu fish were higher in all populations. There was higher citrate synthase activity in LUIB fish than in OBSE or ARDH (Fig. 7; GLM with Type I sums of squares: liver weight F = 0.11 p = 0.740633; population F = 6.66, p = 0.003528; mitochondrial haplotype F = 0.23, p = 0.637131).

**Figure 7.**
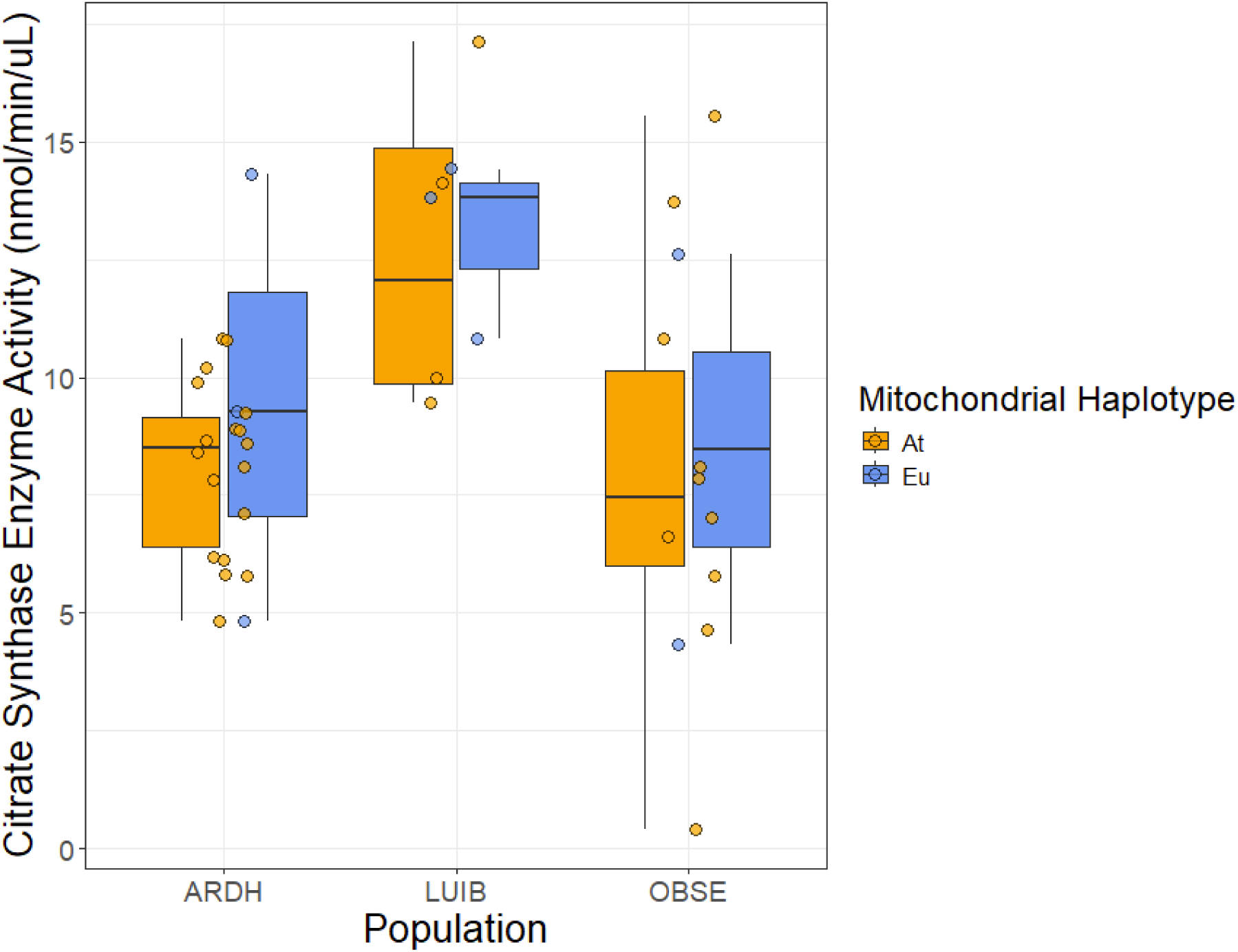
Variation in citrate synthase enzyme activity in the liver of migratory three-spined stickleback from North Uist, between mitochondrial haplotypes and populations (ARDH, LUIB, OBSE). Mitochondrial haplotypes are designated for each population as orange for At and blue for Eu. 25 and 75 quartile ranges are shown as vertical black lines. Horizontal black lines on each box plot represent the median. Each individual point represents one adult stickleback sampled.

## Discussion

Our results show that mitochondrial genetic variation is associated with differences in gonadosomatic index and potentially citrate synthase activity, but overall, differences in traits observed were generally between populations or sexes rather than between haplotypes.

Male GSI was found to be significantly different between mitochondrial haplotypes, with Eu fish having a higher GSI. GSI is a measure of gonad weight relative to body weight and is used to determine reproductive condition. In male fish, GSI is highest during the onset of spermiogenesis, and decreases with the completion of this phase and the elimination of the germ cell cytoplasm. Consequently, a lower GSI could indicate that an individual would have either completed spermiogenesis or is still only in the initial phases of spermatogenesis (Stoumboudi et al. 1997). Given that we observed differences in GSI measurements between haplotypes, it may be that mtDNA influences the timing of male reproductive cycles in stickleback. We know that mitochondria play an important role in spermatogenesis, whereby mitochondria are frequently fragmented and re-built, ending in a sheath surrounding the axoneme (Varuzhanyan et al. 2021). This is when GSI is highest, and this sheath is crucial for sperm mobility (Stoumboudi et al. 1997). Further, mitochondrial energetics are known to influence sperm fertilization capacity and quality, with dysfunctional mitochondria leading to reproductive problems (Ulloa-Rodríguez et al. 2017). Given the complex nature of spermatogenesis and the role of mitochondria in spermiogenesis, it is possible that the underlying mitochondrial genome may affect the timing of these processes. It could be that Eu and At fish exhibit slightly different timescales for mitochondrial dependent processes like mitochondrial fragmentation and sheath formation, therefore affecting the timing of higher or lower GSI measurements.

Another possibility is that the testes of the fish with the Eu haplotype are simply larger, irrespective of reproductive stage. Larger testes are associated with a higher sperm count or potentially bigger sperm cells (Stockley et al. 1996), something that is likely selected based on the environments they are found in, or the morphology of the eggs to be fertilized. We know that abiotic conditions affect egg development on North Uist. Eggs from alkaline lochs hatch less well than eggs from acidic lochs when developed in acidic water, apparently because larvae struggle to escape from the chorion at hatching (Hill 2023). This suggests some difference in chorion make up associated with water chemistry. Given that migratory stickleback on Uist generally breed in quite alkaline conditions and the Eu haplotype is much more common in (especially acidic) freshwater lochs, it is possible that water chemistry has also driven selection for bigger or more sperms cells in Eu fish.

In all populations sampled, Eu fish had higher average hepatic citrate synthase (CS) activity than At fish. Although not significant in the wild-caught fish in this study, we have documented the same pattern in aquarium-raised stickleback, where the difference was statistically significant in these more tightly controlled conditions (Barnes et al. *in review PNAS*). Although CS is encoded by the nuclear genome, its activity is often used as a proxy for mitochondrial content or activity (Larsen et al. 2012) and it is known that feedback from the mitochondria themselves can alter CS activity (Omini et al. 2021, LaNoue et al. 1972). We hypothesise that differences in function of At and Eu genotypes have wider effects on metabolism that alter CS activity at the organismal level in stickleback. Although the functional consequences of specific sequence differences between the Eu and At haplotypes have not been tested, variation in the mitochondrial genome could influence oxidative phosphorylation and mitochondrial signaling, with downstream effects on mitochondrial metabolism and CS activity. The ND4 and ND5 substitutions utilized in our diagnostic assay are synonymous and therefore do not alter the amino acid sequence, however other differences between the haplotypes may affect protein structure and contribute to the observed physiological differences. Given that CS is a commonly used marker of oxidative capacity or mitochondrial density (Larsen et al. 2012), the higher CS activity in Eu fish may reflect increased mitochondrial abundance, potentially in response to differences in mitochondrial efficiency or metabolic requirements between haplotypes. Alternatively, the Eu mitochondria may have a different balance between ATP production and biosynthesis (Lewis et al. *in preparation,* Ishikawa et al. 2019). Given that the Eu lineage is more common and may have evolved in freshwater (Barnes et al. *in review PNAS*) where fatty acids may be in short supply (Ishikawa et al. 2019), we speculate that Eu fish may differ in the allocation of TCA cycle intermediates, including CS, to biosynthesis pathways. This could include fatty acid synthesis or the construction of membrane lipids, although we did not directly investigate these processes. Given that Eu fish exhibited both higher CS activity and higher GSI, differences in mitochondrial metabolism may be associated with variation in reproductive physiology. For example, greater oxidative capacity could potentially support the energetic demands of gonad development, but this relationship, and any underlying mechanisms, remains to be tested.

Body size was bigger in females than males, consistent with stickleback and fish populations globally (Leinonen et al. 2010, Kitano et al. 2012), and is probably because of the fecundity benefits accruing to larger females (Ronco et al. 2019, Horne et al. 2019, Qiao et al. 2016). (Parker 1992). In stickleback, other factors often influence male reproductive success, such as coloration (redder males tending to be more successful in mating), so size is not the only factor driving their sexual success (Bakker & Mundwiler 1994). Differences in length between populations are presumably a result of geographical variation in environmental factors and have been documented in other migratory stickleback populations (Blouw & Hagen 1990, Morris et al. 2018). The fact that this difference between populations persists at a small geographical scale suggests either that migratory stickleback have some degree of local adaptation and therefore must usually return to breed in the population where they were themselves born, or that early life experience has a long-term effect on size. Migratory stickleback spend about three months in their natal waterbody before migrating to sea, so the latter is possible, although it seems unlikely that these early differences would persist after a further eight to nine months feeding at sea, given the propensity of stickleback for catch-up growth (Álvarez & Metcalf 2005, Metcalf & Monaghan 2001).

Female stickleback had larger livers (HSI and liver weight per gram) while male stickleback had larger pectoral muscles and more fat. Sexual dimorphism in fish livers is well documented, as female liver cells contain larger reserve vesicles in addition to increased granular endoplasmic reticulum, mitochondria, and peroxisomes (Qiao et al. 2016). In female fish, the liver plays a major role in oocyte production, the process by which an immature ovarian cell is produced (Qiao et al. 2016), and this alone could explain its larger size in females. Larger pectoral muscles and more fat in males are likely related to stickleback parental care strategies. Male stickleback provide parental care through the continuous “fanning” of oxygenated water over fertilized eggs in their nest and do this for the duration of egg development (Lachance and Fitzgeralds 1992, Bakker et al. 2006). The pectoral muscle is the main muscle used for this “fanning” motion and is likely why the male stickleback have larger pectoral muscles. Additionally, the greater fat reserves of the male stickleback are likely useful when caring for developing eggs. As these males will need to provide constant care, it is likely that their foraging is restricted and they would need to draw on the nutrients stored within their fat reserves.

Nutrient availability may explain discrepancies in liver size between populations, as specific food items and diet can affect biomass gain differently. Alternatively, reproductive timing could also be an explanation for organ size variation, as reproductive cycles in combination with nutrient availability are known to affect body protein and lipid depletion and retention (Talbot 1993). Migratory stickleback populations on North Uist do exhibit variation in the timing of reproduction (A.D.C. MacColl, personal observations). Fish in populations that started breeding earlier would be expected to have lower reserves.

## Conclusion

Migratory stickleback populations on North Uist provide an ideal natural system for studying the consequences of mitochondrial variation (Beck et al. 2022). The two deeply diverged mitochondrial haplotypes segregate against a freely intermixed nuclear background, an unusual situation in wild populations providing a natural equivalent of laboratory conplastic studies. Here we have shown that while most phenotypic traits that we measured do not covary with mitochondrial haplotype, nevertheless male gonadosomatic index and citrate synthase activity both do, suggesting avenues for further research investigating the links between mitochondrial variation, physiology and reproductive traits.

